# Revealing the Oligomerization of Channelrhodopsin-2 in the Cell Membrane using Photo-Activated Localization Microscopy

**DOI:** 10.1101/2023.05.24.542088

**Authors:** Ekaterina Bestsennaia, Ivan Maslov, Taras Balandin, Alexey Alekseev, Anna Yudenko, Assalla Abu Shamseye, Dmitrii Zabelskii, Arnd Baumann, Claudia Catapano, Christos Karathanasis, Valentin Gordeliy, Mike Heilemann, Thomas Gensch, Valentin Borshchevskiy

## Abstract

Microbial rhodopsins are retinal membrane proteins that found a broad application in optogenetics. The oligomeric state of rhodopsins is important for their functionality and stability. Of particular interest is the oligomeric state in the cellular native membrane environment. Fluorescence microscopy provides powerful tools to determine the oligomeric state of membrane proteins directly in cells. Among these methods is quantitative photoactivated localization microscopy (qPALM) allowing the investigation of molecular organization at the level of single protein clusters. Here, we apply qPALM to investigate the oligomeric state of the first and most used optogenetic tool Channelrhodopsin-2 (ChR2) in the plasma membrane of eukaryotic cells. ChR2 appeared predominantly as a dimer in the cell membrane and did not form higher oligomers. The disulfide bonds between Cys34 and Cys36 of adjacent ChR2 monomers were not required for dimer formation and mutations disrupting these bonds resulted in only partial monomerization of ChR2. The monomeric fraction increased when the total concentration of mutant ChR2 in the membrane was low. The dissociation constant was estimated for this partially monomerized mutant ChR2 as 2.2±0.9 proteins/μm^2^. Our findings are important for understanding the mechanistic basis of ChR2 activity as well as for improving existing and developing future optogenetic tools.

## Introduction

Microbial rhodopsins constitute a large group of light-sensitive proteins with seven-transmembrane α-helices found in pro- and eukaryotic microorganisms. These proteins harbor a retinal cofactor which is photoisomerized upon illumination. This process is the primary event underlying the diverse functions of retinal proteins as light-driven pumps, light-gated channels and photoreceptors. Expression of microbial rhodopsins in neurons enables to control the nerve cells’ membrane potential by light with high temporal and spatial resolution ^[1]^. This became the basis for the emergence and development of a technique called optogenetics ^[2]^. It allows using light to control and study cellular activities in isolated neurons as well as complex neural systems such as the brain of living animals ^[3] [4]^. Promising biomedical applications of rhodopsins in optogenetics are aimed to restore vision ^[5]^, hearing ^[6]^ and memory ^[7]^.

Microbial rhodopsins can form stable oligomers of various stoichiometry (from dimers to hexamers) ^[8]^ that are relevant to their properties and function. The best-known example is bacteriorhodopsin (BR) – the archetypal light-driven proton pump of *Halobacterium salinarum*. BR forms trimers packed into a two-dimensional lattice in the native purple membrane ^[9]^. The trimeric form of BR has significantly higher proton pumping efficiency ^[10]^ and is important for its higher thermal and photostability ^[11–13]^ relative to the monomeric state. Other examples of oligomerization-dependent functions include Na^+^-pumping by pentametric *Krokinobacter* eikastus rhodopsin 2 (KR2) ^[14]^, proton pumping by hexameric proteorhodopsin (PR)^[15,16]^ and ion channeling by trimeric ChRmine ^[17]^ or pentameric Organic Lake Phycodnavirus rhodopsin II (OLPVRII)^[16]^. In most cases, the oligomeric state of microbial rhodopsins was determined by atomic force microscopy ^[18,19]^, circular dichroism spectroscopy ^[19–21]^, electron microscopy ^[22,23]^, crosslinking ^[15,16]^ and X-ray crystallography ^[24–26]^. These methods require sophisticated sample preparation that in many cases may imprint a non-natural oligomeric state to the microbial rhodopsins under study^[8]^. Hence, although numerous new microbial rhodopsins from various organisms were recently identified, the information about their oligomeric states remained sparse and largely incomplete because of a lack of approaches to determine their oligomeric state in a native cell environment.

Channelrhodopsin-2 (ChR2) is a light-gated cation channel found in the unicellular green alga *Chlamydomonas reinhardtii* ^[27]^. It is the first^[1]^ and currently the most widely used rhodopsin in optogenetics^[2]^. Mass spectrometry of protein detergent solution confirmed by electron crystallography of 2D-crystals showed that ChR2 forms dimers ^[23,28,29]^. These findings, however, do not allow concluding unequivocally that the dimer is ChR2’s functional unit, since in both sample preparations ChR2 is placed in a non-natural environment. The X-ray diffraction crystal structure of ChR2 contains two symmetrical interprotomer disulfide bridges formed by Cys34 and Cys36^[26]^. Such N-terminal extracellular cysteine residues are conserved in other structurally characterized dimeric channelrhodopsins ^[30–32]^. Mutations of Cys34/Cys36 do not destroy ChR2 dimers when purified protein samples were analyzed by EPR ^[33,34]^ and no effect on ChR2-induced photocurrents was detected in ChR2-expressing oocytes ^[35]^. The oligomeric state of ChR2 and its mutants have not been investigated in the plasma membrane of a cell so far and therefore the nature of its functional unit remains unclear.

Fluorescence microscopy provides a variety of methods to analyze membrane protein oligomerization in an intact cell environment. These methods include Förster resonance electron transfer (FRET) ^[36]^, number & brightness analysis (N&B) ^[37,38]^, spatial intensity distribution analysis (SpIDA) ^[39,40]^, fluorescence correlation spectroscopy (FCS) ^[41,42]^, stepwise fluorescence photobleaching ^[43,44]^, single-molecule tracking ^[45,46]^, point accumulation in nanoscale topography (PAINT) ^[47,48]^, and photoactivated localization microscopy (PALM) ^[49,50]^.

Typical cell surface density of ChR2 used in optogenetics can be estimated from photocurrent noise in patch clamp experiment as ≈200 proteins/μm^2 [51]^ that also corresponds to those obtained by direct counting on the freeze-fracture electron microscopy images ^[52]^. This high density of protein does not allow to spatially resolve individual ChR2 oligomers via diffraction-limited fluorescence microscopy. Single-molecule localization microscopy (SMLM) ^[53]^ overcomes this resolution barrier, and in addition to spatially resolving single protein clusters in intact cells, provides information on molecule numbers and protein oligomerization within single protein clusters ^[54]^. Here, we use SMLM in combination with photoswitchable fluorescent proteins (PALM), where sparsity of active emitters is controlled via photoconversion and photobleaching of fluorescent proteins genetically fused to the protein of interest (Figure 1a). We use a kinetics-based analysis, quantitative PALM (qPALM), which relates the number of single-molecule emission events to molecule numbers ^[55]^ (Figure 1b,c,d) and informs on protein oligomerization ^[56]^. The accurate analysis takes into account two challenges when using fluorescent proteins: first, some fluorescent proteins are not detected due to e.g. incomplete maturation, which results in undercounting; second, some fluorescent proteins produce more than one fluorescent burst, due to reversible transition into long-lived non-fluorescent states, which results in overcounting ^[57]^. Quantitative PALM analysis inherently corrects for both incomplete detection and multiple emission events^[55]^. In a best-practice procedure, the experimental parameters describing the detection efficiency and the degree of multi-signal detection are determined from experiments with reference proteins with known oligomeric states^[50]^.

**Figure 1.**
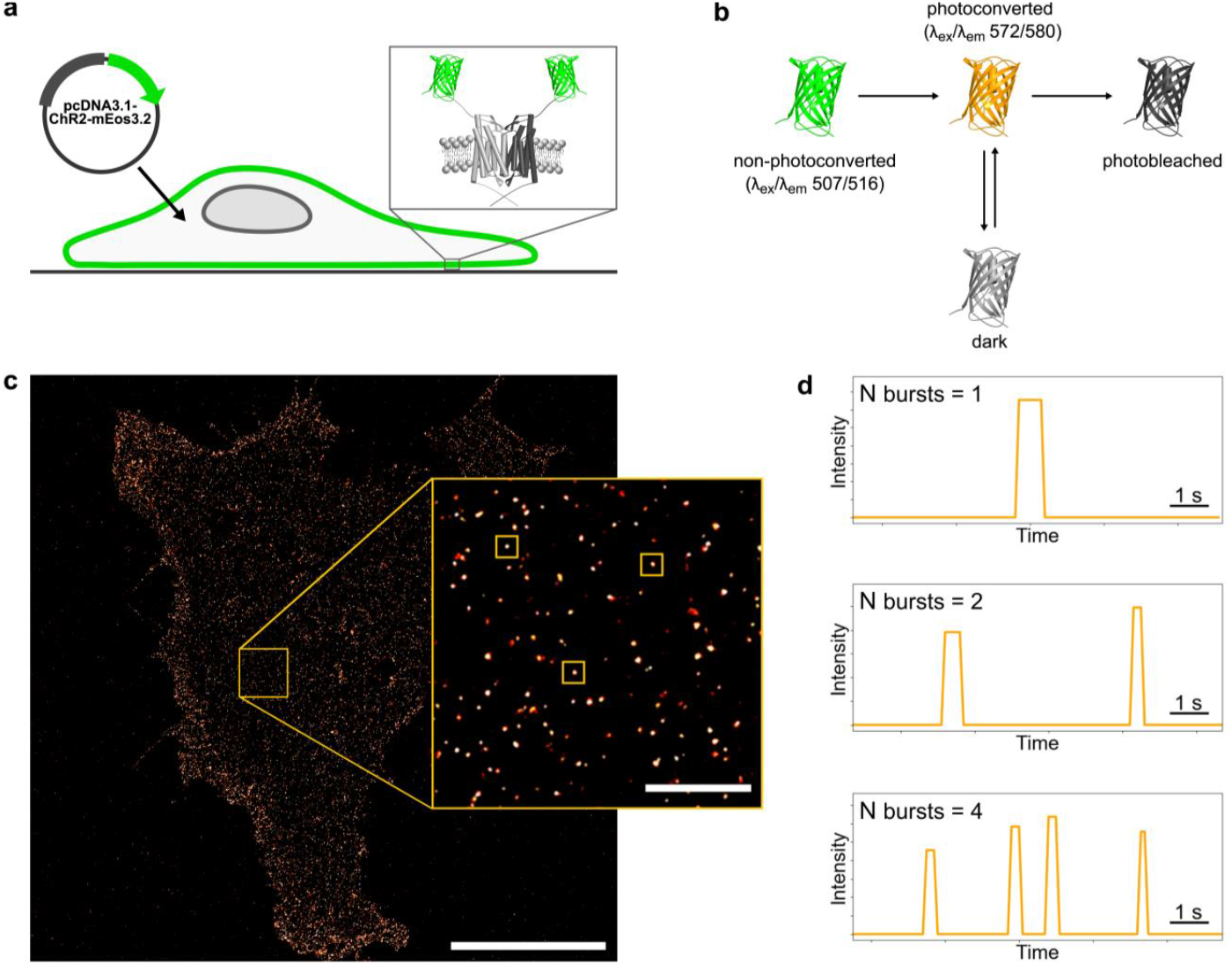
Illustration of general principles underlying quantitative PALM experiment. **(a)** HEK293 cells were transiently transfected with a pcDNA3.1 plasmid coding ChR2 fused to mEos3.2. The fluorescence from proteins in the basal plasma membrane was analyzed via quantitative PALM. **(b)** 4-state photokinetics model ^[83]^ describing photoconvertible fluorescent protein mEos3.2. During PALM experiment mEos3.2 first is photoconverted by violet light (405 nm) from a shorter wavelength (non-photoconverted form; λ_ex_/λ_em_ (nm) 507/516) to a longer wavelength emission state (photoconverted form; λ_ex_/λ_em_ (nm) 572/580). Photoconverted mEos3.2 can photoblink, i.e. reversibly switch between a non-emissive dark state and the emissive photoconverted state until irreversible photobleaching.**(c)** An example of a super-resolved cell membrane image obtained from a PALM experiment. PALM experiments are performed on cells expressing membrane proteins fused with mEos3.2 (ChR2_WT_-mEos3.2 in this particular case). The fluorescence of a single photoconverted mEos3.2 is detected in TIRF mode allowing high-precision visualization of the proteins localized in the basal plasma membrane; scale bar 10 μm. Zoomed inset: individual protein clusters can be selected and the number of fluorescent bursts in individual clusters can be determined; scale bar 1 μm. **(d)** Examples of fluorescence time courses in individual clusters revealing the number of bursts.

In this work, we determined the oligomeric state of ChR2 in the intact plasma membrane of fixed human embryonic kidney (HEK293) cells using qPALM. We compare ChR2 with well-characterized monomeric and dimeric membrane protein reference samples and show that wild-type ChR2 (ChR2_WT_) predominantly forms dimers in the plasma membrane. Furthermore, we analyze the role of two N-terminal cysteines Cys34 and Cys36 that stabilize the ChR2 dimer via disulfide bonds. We show that mutation of these cysteines partially decreases the fraction of ChR2 dimers while simultaneously increasing the fraction of ChR2 monomers. The extent of this effect depends on the ChR2 density in the plasma membrane that allows us to determine equilibrium dissociation constant of dimerization.

## Results

Membrane proteins with known oligomeric states were used to calibrate ChR2 molecular counting. As references, we chose monomeric β_1_-adrenergic receptor (β_1_AR) and constitutively dimeric CD28, which have been used as standard controls in previous studies ^[58,59]^. Both proteins were tagged with photoconvertible fluorescent protein mEos3.2. It is a brighter and truly monomeric version of mEos2 ^[60]^ successfully used in other qPALM-based studies ^[61]^. Both reference constructs β_1_AR-mEos3.2 and CD28-mEos3.2 were expressed in human embryonic kidney 293 (HEK293) cells and mostly localized in the plasma membrane (Figure 2a). The HEK293 cell line was chosen as a common system for optogenetic functional tests of microbial rhodopsins ^[31,62]^. Additionally, this cell line does not express either β_1_AR or CD28 endogenously ^[63]^. We used PALM images to visually separate individual protein oligomers on the cell membrane. On super-resolved images obtained from PALM, individual oligomers appear as bright round-shaped clusters of less than 100 nm in size (Figure 1c). We counted the number of fluorescent bursts of mEos3.2 in individual clusters. As expected, the clusters of dimeric CD28 contain more fluorescent bursts than monomeric β_1_AR: the most frequent numbers of bursts in a cluster are two and one for CD28 and β_1_AR, respectively (Figure 2b,c).

**Figure 2.**
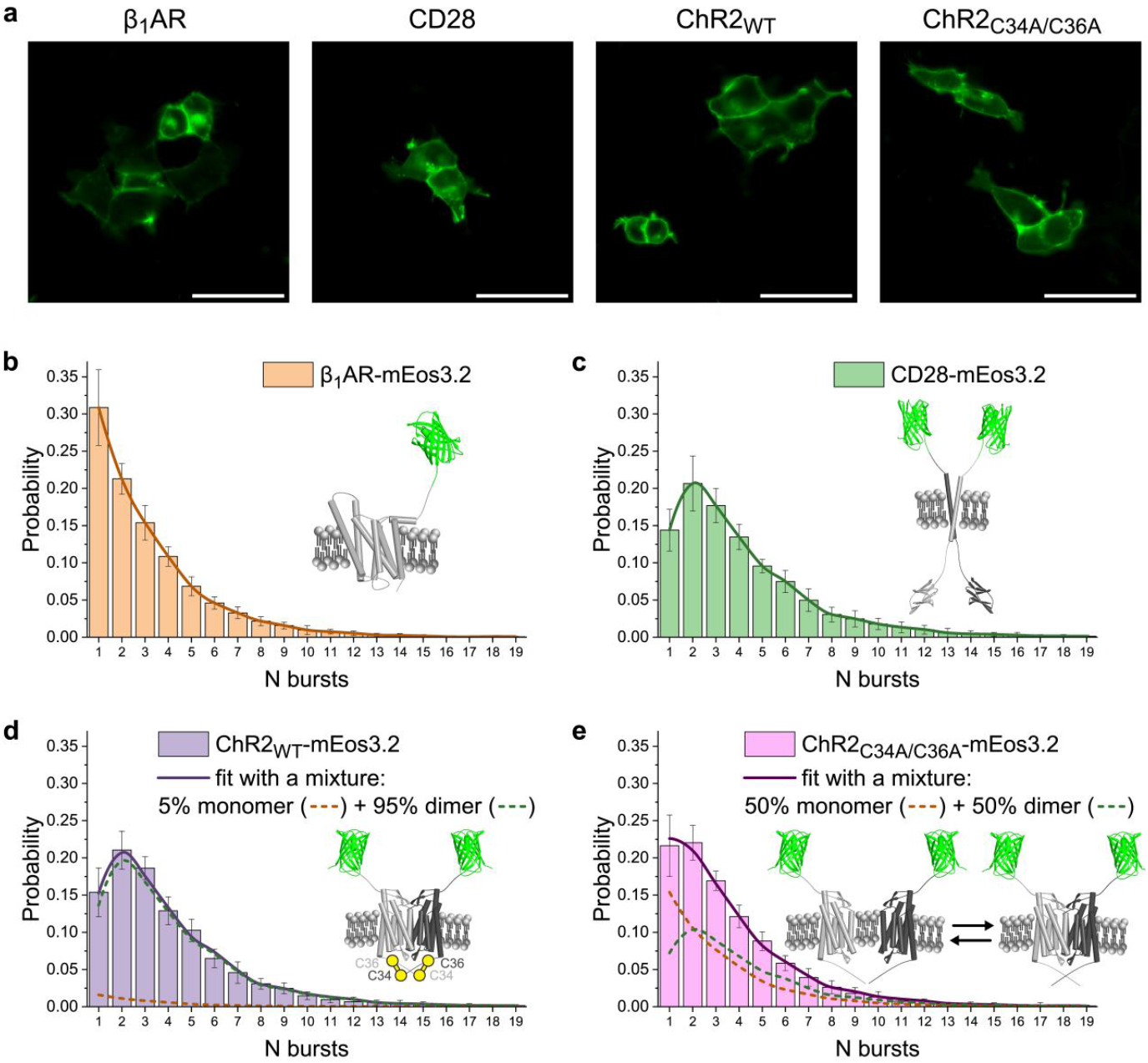
Confocal images of transfected cells and burst distributions for four proteins used in the study: β_1_AR, CD28, ChR2_WT_, and ChR2_C34A/C36A_. **(a)** Confocal images of HEK293 cells expressing membrane proteins fused with mEos3.2. The green form of mEos3.2 was excited with an argon ion laser at 488 nm. Scale bar 50 μm. **(b**,**c**,**d**,**e)** Numbers of fluorescent bursts in individual clusters of membrane proteins fused with mEos3.2. Distributions were averaged over individual cells. Error bars represent the standard deviation. **(b**,**c)** Burst distributions obtained for β_1_AR-mEos3.2 (14 cells, 10,631 clusters) **(b)** and CD28-mEos3.2 (15 cells, 7,844 clusters) **(c)**, which were used as monomeric and dimeric references, respectively. **(d)** Burst distribution obtained for ChR2_WT_-mEos3.2 (12 cells, 7,510 clusters) was fitted to the weighted sum of reference distributions (weighted components are indicated by dashed lines). ChR2_WT_ showed predominant dimerization (95% of dimers) in the cell membrane. **(e)** Burst distribution obtained for ChR2_C34A/C36A_-mEos3.2 (11 cells, 6,966 clusters) was fitted to the weighted sum of reference distributions (weighted components are indicated by dashed lines). The mutation of the two cysteines caused the partial decrease of ChR2 dimer fraction accompanied by the increase of monomer fraction (from 5% of monomers to 50% of monomers).

To determine the oligomerization of ChR2_WT_ in the plasma membrane we transfected HEK293 cells with a ChR2_WT_-mEos3.2 construct. ChR2_WT_ was predominantly localized in the plasma membrane as it was the case for the reference membrane proteins β_1_AR and CD28 (Figure 2a). We visualized individual ChR2_WT_ clusters in the plasma membrane using PALM and counted the number of mEos3.2 fluorescent bursts in individual clusters (Figure 2d). The histogram of fluorescent bursts in individual clusters for ChR2_WT_ appears very similar to the burst distribution of the dimer reference CD28. ChR2_WT_ burst distribution was well fitted by the weighted sum of distributions pre-recorded for monomeric β_1_AR and dimeric CD28, showing no evidence of higher oligomerization (no right shift of the distribution maximum compared to the distribution of dimeric CD28). According to this approximation, ChR2_WT_ is almost fully dimerized in the cell membrane (95% of dimers and 5% of monomers). To estimate the significance of this approximation, we evaluated cell-to-cell variations. The data from individual cells were processed in the same manner (fitted by the weighted sum of reference distributions; Figures S1-S3). Despite slight variations in the dimer fraction between individual cells, statistical analysis showed no significant difference between ChR2_WT_ and reference dimer CD28 (Figure 3a).

**Figure 3.**
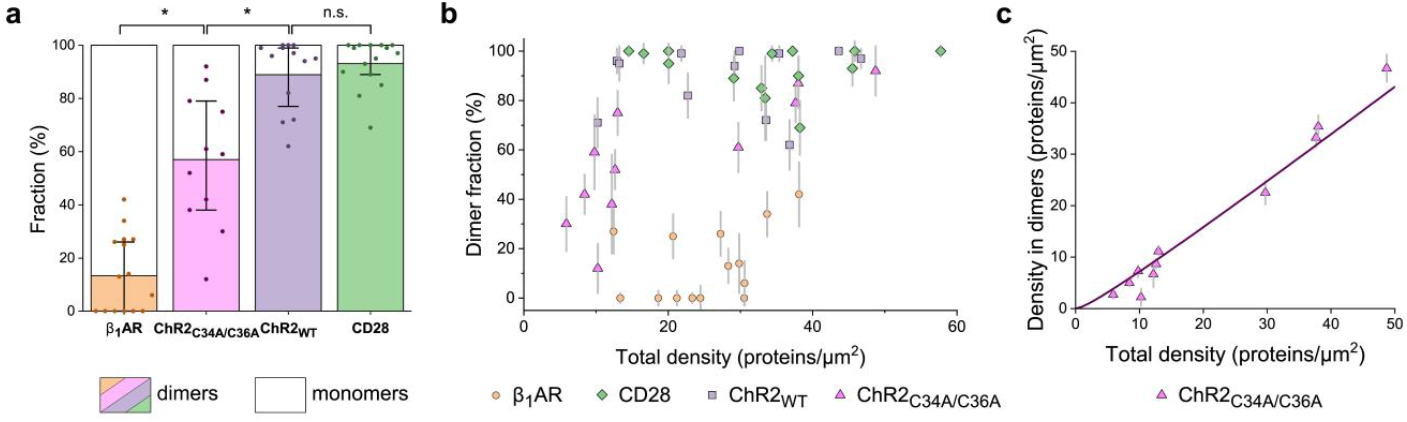
Analysis of cell-to-cell variations in PALM measurements. **(a)** Distributions of numbers of fluorescent bursts in individual clusters obtained from individual cells were fitted with the weighted sum of reference monomeric and dimeric distributions. Fractions of monomer and dimer clusters in single cells are shown as individual data points; the mean and the inter-quartile range are shown as bars with whiskers. Taking into account deviations caused by poorer statistics available from individual cells and their non-ideal homogeneity, ChR2_WT_ does not significantly differ from reference dimer CD28. At the same time, ChR2_C34A/C36A_ showed the widest cell heterogeneity and on average significantly differed both from reference monomer β_1_AR and ChR2_WT_; * p < 0.05 (Kruskal-Wallis test with post-hoc Dunn’s test). **(b)** Dependence between the fraction of dimers and average protein density in the plasma membrane. Protein density is proportional to the number of bursts per unit area in the PALM experiment. From burst distributions for β_1_AR we estimated that one protein emits on average 3.25 bursts. Every point represents one cell expressing monomeric β_1_AR (orange circles), dimeric CD28 (green rhombuses), ChR2_WT_ (purple squares) or ChR2_C34A/C36A_ (pink triangles). Errors are estimated with a bootstrapping algorithm. As well as reference proteins, ChR2_WT_ demonstrates no clear tendency to change oligomers ratio depending on protein density in the plasma membrane. The mutant ChR2_C34A/C36A_ is likely to increase dimer fraction with increasing protein density. **(c)** For ChR2_C34A/C36A,_ the dependence of density of proteins residing in dimers on the total density of proteins is fitted to the theoretical formula and the dissociation constant for ChR2_C34A/C36A_ dimers (*K*_*D*_ = 2.2±0.9 proteins/μm^2^) is determined from the fit (solid line; see “Determination of dissociation constant *K*_*D*_ for ChR2_C34A/C36A_ dimers” in Materials and Methods).

In a crystal structure^[26]^, the ChR2 dimer is stabilized by two covalent interprotein bonds between cysteines at positions 34 and 36. To test the effect of these bonds on the oligomeric state, we prepared a double-mutant ChR2 version (ChR2_C34A/C36A_), tagged with mEos3.2. The mutations of ChR2 did not affect the predominant localization of the protein in the plasma membrane (Figure 2a). As before, we counted the number of bursts in individual ChR2_C34A/C36A_ clusters visualized with PALM. The burst distribution averaged across all measured cells showed a larger fraction of 1-burst clusters for ChR2_C34A/C36A_ compared to ChR2_WT_ (Figure 2d, e). Again, the distribution for the number of fluorescent bursts was fitted to the weighted sum of reference monomeric and dimeric distributions (Figure 2e). The dimeric fraction of ChR2 decreased from 95% in ChR2_WT_ to 50% in ChR2_C34A/C36A_, while the monomer fraction increased concomitantly from 5% to 50%. The significance of this change in monomer-dimer equilibrium was estimated by evaluating cell-to-cell variations in our PALM measurements. To calculate the fraction of dimeric ChR2_C34A/C36A_ in single cells we fitted the individual burst distributions per cell with the weighted sum of reference distributions (Figure S4, Figure 3a). The fraction of dimers was significantly lower in ChR2_C34A/C36A_ than in ChR2_WT_ (p < 0.05).

In comparison to ChR2_WT_, the mutant ChR2C34A/C36A showed higher cell-to-cell variability of the dimeric fraction ranging from 12% to 92% in individual cells (Figure 3a). We proposed that in the absence of stabilizing disulfide bonds, dimerization of ChR2_C34A/C36A_ is transient and the fraction of dimers varies depending on the density of proteins per unit area of the plasma membrane. To test this hypothesis, we looked at the dependence of the dimer fraction on the density of proteins in the plasma membrane (Figure 3b). Since membrane proteins are stoichiometrically labeled, the density of fluorescence bursts on the membrane images is proportional to the protein density. Each protein emits, on average, 3.25 fluorescence bursts, as can be inferred from the blinking distribution of a reference monomer β_1_AR (Figure 2b). The mutant ChR2C34A/C36A showed a statistically significant trend from predominant monomeric fraction at low protein densities towards predominant dimeric fraction at higher densities (Spearman’s rank correlation coefficient *r* = -0.80, *p* = 0.003). On the contrary, there was no statistically significant correlation between the monomeric fraction and protein density for ChR2_WT_ or reference proteins (*p* > 0.1). Finally, we determined the dissociation constant for ChR2_C34A/C36A_ dimers via fitting the dependence between density of proteins residing in dimers and total density of proteins with a theoretical curve (*K*_*D*_ = 2.2±0.9 proteins/μm^2^, Figure 3c, see “Determination of dissociation constant *K*_*D*_ for ChR2_C34A/C36A_ dimers” in Materials and Methods).

## Discussion

In this work, we applied qPALM to determine the degree of ChR2 oligomerization directly in the cellular plasma membrane. ChR2_WT_ showed no significant difference from the reference dimeric protein CD28. This confirms that most of the protein in cell membrane forms dimers as it was expected from previous studies with purified ChR2, where stable dimers were observed via crystallography ^[23,26,29]^, mass-spectrometry^[28]^ and gel electrophoresis ^[23,28]^. While these earlier results pointed towards a strong dimerization tendency, the formation of higher ChR2 oligomers in the plasma membrane could not be excluded. The absence of higher oligomers in the cell membrane was confirmed by our qPALM experiments.

We also tested the contribution of inter-domain disulfide bonds between Cys34 and Cys36 to the formation of ChR2 dimers. In contrast to the stable dimerization of ChR2_WT_, the mutant ChR2_C34A/C36A_ was, on average, a mixture of 50% monomers and 50% dimers in the cell membrane. This result suggests that inter-protomer disulfide bonds are important but not necessary for the dimer formation. In the previous EPR-based studies, purified ChR2 remained dimeric even when inter-protomer disulfide bonds were removed ^[33,34,64]^. Based on the crystallographic structure^[26]^, it can be inferred that the dimer is stabilized through a combination of backbone H-bonds, specific interactions between side chains as well as several water-mediated interactions. The higher fraction of dimers in purified ChR2 compared to our study can be explained by high density of the protein due to overexpression in a heterologous expression system, altered lipid environment after solubilization and high protein concentrations required for EPR measurements.

It is particularly intriguing whether the monomeric form of the ChR2_C34A/C36A_ is functional. Structural^[26]^ and mutagenesis ^[35]^ studies of ChR2 showed that the monomer alone contains all necessary structural elements (gates and cavities) for ion channeling. It is therefore generally accepted that a monomer rather than a dimer is a functional unit of ChR2. However, even for the functional monomer, oligomerization may be a necessary requirement (similar to ChRmine^[17]^) or at least significantly enhance its efficiency (similar to BR^[10]^). Oligomerization might well be important for operative orientation of monomers in the lipid bilayer and stabilize their structural elements in functional states as it was shown for KR2 ^[65]^. Although previous studies showed that ChR2 without inter-subunit disulfide bonds retains photocurrent activity in oocytes ^[35,64]^, the accurate correlative analysis of the ChR2 activity, density and oligomerization will be required to determine whether ChR2 monomers are functional.

Without stabilizing inter-subunit disulfide bonds, the ratio between monomers and dimers in ChR2_C34A/C36A_ shifts towards pure monomers at low total protein densities in the membrane. This anti-correlation between dimeric fraction and protein density suggests that a dynamic balance exists between monomers and transient dimers in ChR2_C34A/C36A_. The dissociation constant for ChR2_C34A/C36A_ dimers (*K*_*D*_) can be estimated as 2.2±0.9 proteins/μm^2^. This finding implies that mutant ChR2_C34/C36_ must be predominantly dimeric at the expression levels common in optogenetic applications (∼200 proteins/μm^2, [51]^).

The estimated dissociation constant for ChR2_C34A/C36A_ dimers is similar to those previously reported for other transiently dimerizing seven transmembrane helix proteins, class A G-protein coupled receptors (GPCRs) ^[66–71]^. In GPCRs, dimerization is tightly coupled to the functioning of receptors. Dimerization impacts the trafficking of receptors to the plasma membrane, regulates receptor internalization ^[72]^, affects ligand binding and downstream signaling ^[73,74]^. On the other side, the equilibrium between monomers and dimers depends on the activation state of the proteins ^[59,69]^. Given the structural similarity between retinal proteins and class A GPCRs and similarity between their dimer dissociation constants, it is reasonable to expect similar effects of dimerization for retinal proteins. Indeed, oligomerization was proposed to hinder the development of fused tandem rhodopsins due to undesired clustering of proteins ^[75]^. Multimerization properties were also suspected to completely deteriorate channelrhodopsin-derived tandem rhodopsins function in lysosomes ^[76]^. These findings highlight the importance of understanding the effects of dimerization and oligomerization when developing optogenetic tools.

Our study demonstrates the successful application of qPALM to determine the oligomeric state of rhodopsins in intact cells. This proof-of-concept study paves the way for further qPALM-based investigations of rhodopsins’ oligomerization. In combination with functional tests, qPALM is potentially useful for mechanism-of-action studies that can reveal whether minimal functional units of specific rhodopsins are monomers or oligomers. Future qPALM studies can facilitate engineering of optogenetic tools whose functionality is affected by rhodopsin oligomerization.

## Materials and methods

### Plasmids

The gene of C-terminally truncated ChR2_WT_ fused with yellow fluorescent protein (EYFP) ^[51]^ was kindly provided by Prof. E. Bamberg (Max Planck Institute of Biophysics, Frankfurt, Germany) and inserted into pcDNA3.1 plasmid. Then the gene for EYFP was replaced with the gene for mEos3.2 used previously ^[61]^.

For expression of ChR2_C34A/C36A_-mEos3.2 both mutations C34A and C36A were introduced in pcDNA3.1-ChR2_WT_-mEos3.2 plasmid with one pair of partially complementary primers in a single PCR ^[77]^.

The plasmid pcDNA3-β_1_AR-EYFP was a gift by the group of P. Annibale & M.J. Lohse (Max Delbrück Center, Berlin, Germany). The gene for EYFP was replaced with the gene for mEos3.2 using XbaI and ApaI restriction sites.

CD28 gene (truncated at R185) was amplified from a commercially available plasmid (pEF6a-CD28-PafA, Addgene, #113400) and inserted into the plasmid pcDNA3.1-ChR2_WT_-mEos3.2 instead of ChR2 gene using BamHI and NotI restriction sites.

The full DNA sequences of the used constructs are provided in Table S1.

### Cell culture and sample preparation for microscopy

Flp-In™ T-REx™ 293 cells (referred to as HEK293 through the manuscript; Thermo Fisher Scientific, Waltham, Massachusetts, USA) were maintained at 37 °C in a 5% CO_2_ humidified atmosphere in DMEM (with high glucose and GlutaMAX™, (Thermo Fisher Scientific, Waltham, Massachusetts, USA) supplemented with 10% fetal bovine serum (Thermo Fisher Scientific, Waltham, Massachusetts, USA) and 50 U/mL Penicillin-Streptomycin (Thermo Fisher Scientific, Waltham, Massachusetts, USA).

HEK293 cells were transfected with 100-1000 ng of plasmid DNA (optimized for each construct) by a modified calcium-phosphate method ^[78]^ in 6 cm cell culture dishes. In the case of transfections with ChR2 variants, 1 μM all-trans retinal was added to the cell medium to ensure sufficient chromophore for proper maturation of the rhodopsin. Cells were kept at 37°C, 5% CO_2_, and in a humidified atmosphere. One day after transfection, cells were seeded on 9 mm diameter glass coverslips for confocal fluorescence microscopy or in 35 mm imaging dishes with a glass bottom (ibidi) for PALM and kept for another 24h at 37°C, 5% CO_2_ in a humidified atmosphere. All glass surfaces were pre-coated with poly-L-lysine (0.1 mg/mL; Merck, Dramstadt, Germany). Transfected cells were carefully washed with PBS several times and fixed with 4% paraformaldehyde solution in PBS for 30 min at room temperature. After fixation, samples were repeatedly washed with PBS. PBS was also used as an imaging buffer.

### Confocal microscopy

The localization of the expressed protein was studied using a Zeiss LSM880 laser scanning confocal microscope. Fixed cells were imaged on glass coverslips using Zeiss 20× water immersion objective (NA 1.0). The green form of mEos3.2 was excited with an argon-ion laser at 488 nm and fluorescent emission was detected after passing a suitable emission filter (499-541 nm) using a GaAsP photomultiplier tube.

### Single-molecule localization microscopy

PALM was performed on a custom-built setup described previously ^[79]^. It is based on an Olympus IX-71 inverted microscope body equipped with an Olympus ApoN 60× oil objective (NA 1.49). Fixed cells were imaged in 35 mm glass bottom microscopy dishes. To find transfected cells with sufficient localization of the protein in the membrane, we excited the green form of mEos3.2 in wide-field mode at 488 nm (<1 mW laser power at the sample, Ar-ion laser, Innova 70C; Coherent). PALM movies were recorded in total internal reflection fluorescence (TIRF) mode. For PALM, mEos3.2 was simultaneously photoconverted, imaged and photobleached by gradually increasing UV illumination (405 nm; up to 1 mW at the sample; Cube 405-100C; Coherent) combined with continuous excitation at 561 nm (approximately 50 mW at the sample; Sapphire 561-200 CDRH-CP; Coherent). Image series (6,000 - 12,000 frames with 100 ms acquisition time per frame) were recorded using an EMCCD camera (AndoriXon DU897E; Andor) with 512 × 512 pixels and a pixel size of 80 nm.

### Data analysis

To define single-molecule localizations and create super-resolved images of the cell membrane, PALM movies were processed in rapi*d*STORM software ^[80]^. In brief, single mEos3.2 fluorescent bursts were localized in each frame using two-dimensional Gaussian PSF fitting with PSF FWHM of 270 nm and signal-to-noise threshold of 9. Localizations identified in consecutive frames were merged. Additionally, one-frame-lasting fluorescent bursts (fluorescence detected at one spatial position in a single frame surrounded by no fluorescence at the preceding and succeeding frames) and one-frame-lasting blinks (no fluorescence detected at one spatial position in a single frame surrounded by fluorescence at the preceding and succeeding frames) were neglected. The final list of localizations was used to generate super-resolved images with a pixel size of 10 nm. Individual protein clusters in super-resolved images were selected manually after a visual inspection in Fiji ^[81]^. The criteria for selecting clusters were their clear separation, high brightness, characteristic size (maximum diameter of 100 nm) and round shape with a single intensity center. The number of fluorescent bursts in selected clusters was calculated with LocAlization Microscopy Analyzer (LAMA ^[82]^). Further analysis of the distributions for numbers of fluorescent bursts in clusters was performed using OriginPro 8.1 or home-written python scripts.

### Determination of dissociation constant K_D_ for ChR2_C34A/C36A_ dimers

To determine ChR2_C34A/C36A_ dimer–monomer equilibrium dissociation constant *K*_*D*_, we fit the dependence of the density of proteins residing in dimers on the total density of the protein in the plasma membrane (Figure 3c). We use the equation for equilibrium dissociation constant of homo-dimerization:

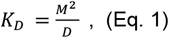

where *M* and *D* are densities of monomeric and dimeric clusters per unit area of the plasma membrane. The total density of the protein in the membrane is:

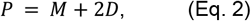

and the fraction of dimeric clusters is

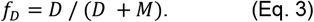

From formulas (2) and (3) we get the expression for the density of proteins residing in dimers 2*D*:

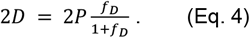

Using equation (4) we calculate the density of proteins residing in dimers 2*D* for each experimental data point shown at Figure 3b. The experimental errors in the fraction of dimeric clusters *Δf*_*D*_ translate into error in 2*D*:

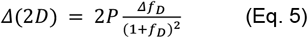

Density of protein molecules in dimers 2*D* follows from (1) and (2):

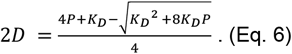

To determine *K*_*D*_, we fitted the experimental data at Fig. 3c according to the equation (6) using OriginPro 8.1. To account for experimental errors, weighting factors *ω*_*i*_ = 1/*Δ*(2*D*)^2^ were used for the fitting. The error of the determined *K*_*D*_ was estimated as the standard error of the fit.

## Author contributions

E.B., A.Y., and D.Z. performed the cloning under supervision of T.B.

E.B. and A.A.S. maintained the cell cultures, performed transfections and sample preparation for fluorescence microscopy under supervision of A.B. and T.G.

E.B. performed fluorescence imaging via confocal and super-resolution microscopy under supervision of T.G.

E.B., I.M., C.C., and C.K. processed and analyzed qPALM data under supervision of M.H., T.G. and V.B.

E.B., I.M., T.G., and V.B. prepared a draft of the manuscript.

E.B., I.M., A.A., V.G., M.H., T.G., and V.B. discussed the data, analysis, and contributed to writing the manuscript.

E.B., I.M., T.G., and V.B. conceived the study.

T.G. and V.B. supervised the work.

All authors discussed the data and contributed to editing the manuscript.

## Acknowledgments

We are thankful to Fedor Tsybrov for the help with the preparation of plasmids. V.B. acknowledges DAAD Young Talents Programme Line A. V.G. acknowledges his HGF Professorship. C.C., C.K. and M.H. gratefully acknowledge the Deutsche Forschungsgemeinschaft (grants CRC1507 and CRC807) for financial support.

