## Supplementary material for "Revealing the Oligomerization of Channelrhodopsin-2 in the Cell Membrane using Photo-Activated Localization Microscopy": SI

**Figure S1.** Burst distributions in clusters collected from individual cells expressing $\beta_1$AR-mEos3.2. Each distribution was fitted with the weighted sum of reference monomeric and dimeric distributions, calculated dimer fraction is indicated.

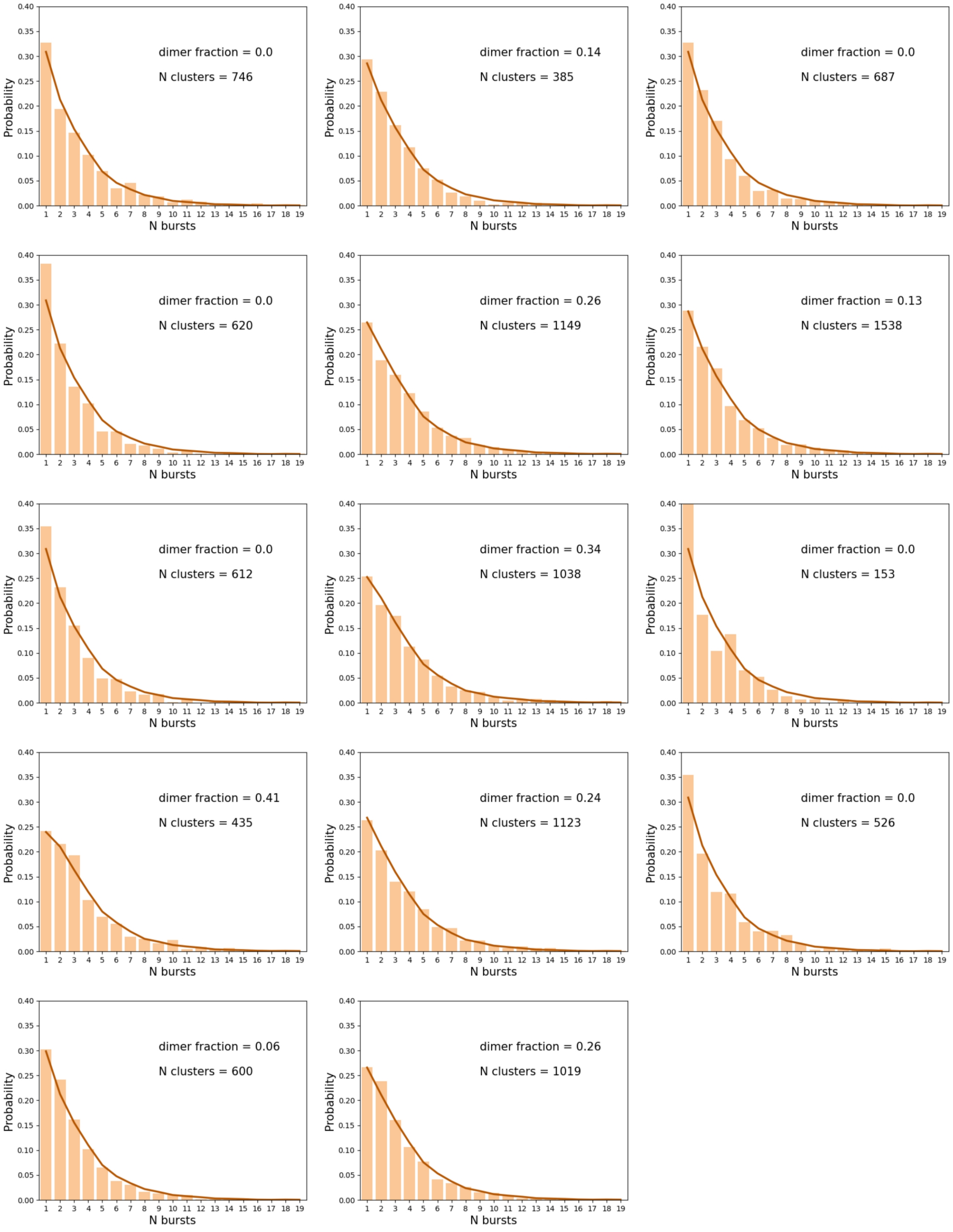

**Figure S2.** Burst distributions in clusters collected from individual cells expressing CD28-mEos3.2. Each distribution was fitted with the weighted sum of reference monomeric and dimeric distributions, calculated dimer fraction is indicated.

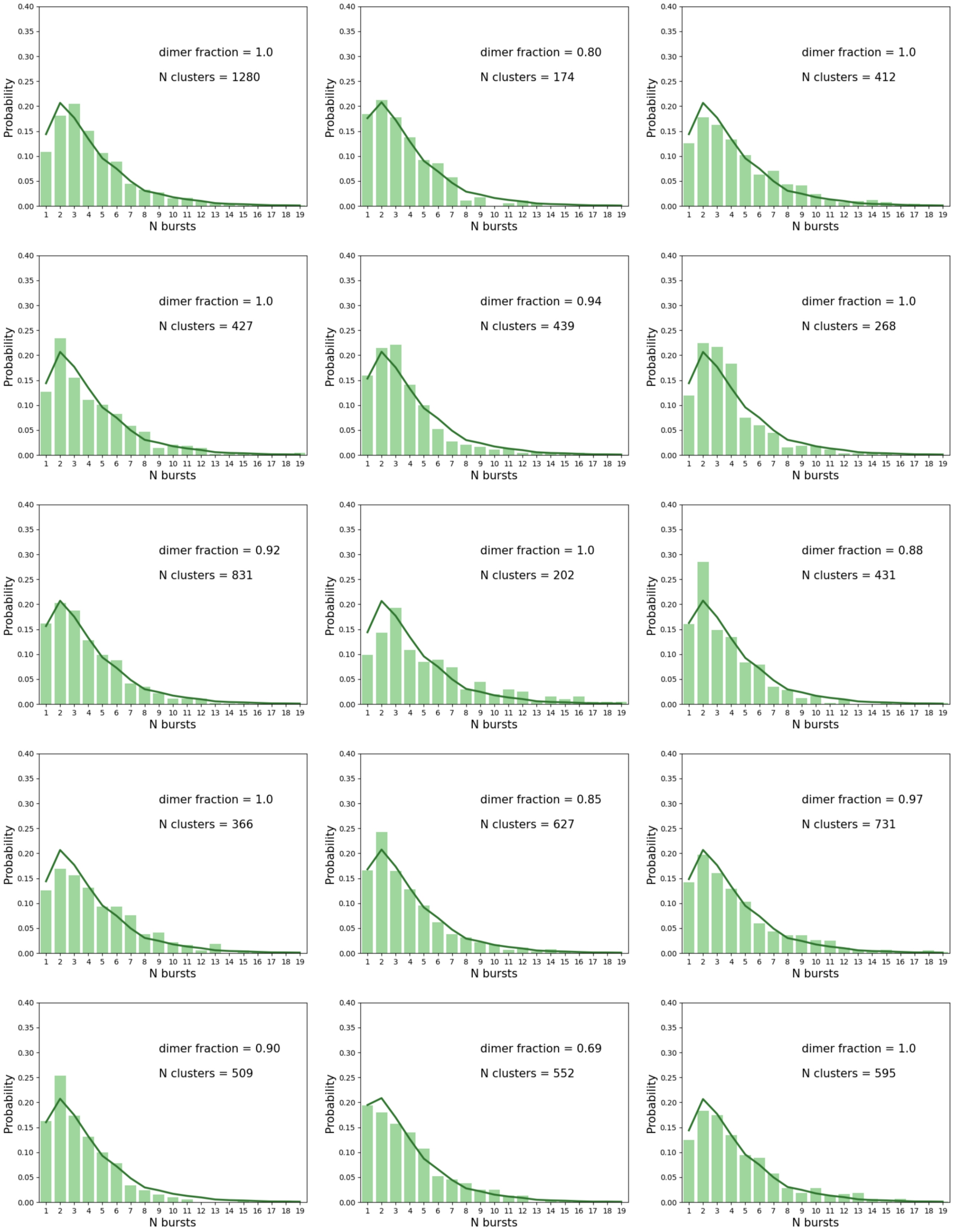

**Figure S3.** Burst distributions in clusters collected from individual cells expressing ChR2$_{WT}$-mEos3.2. Each distribution was fitted with the weighted sum of reference monomeric and dimeric distributions, calculated dimer fraction is indicated.

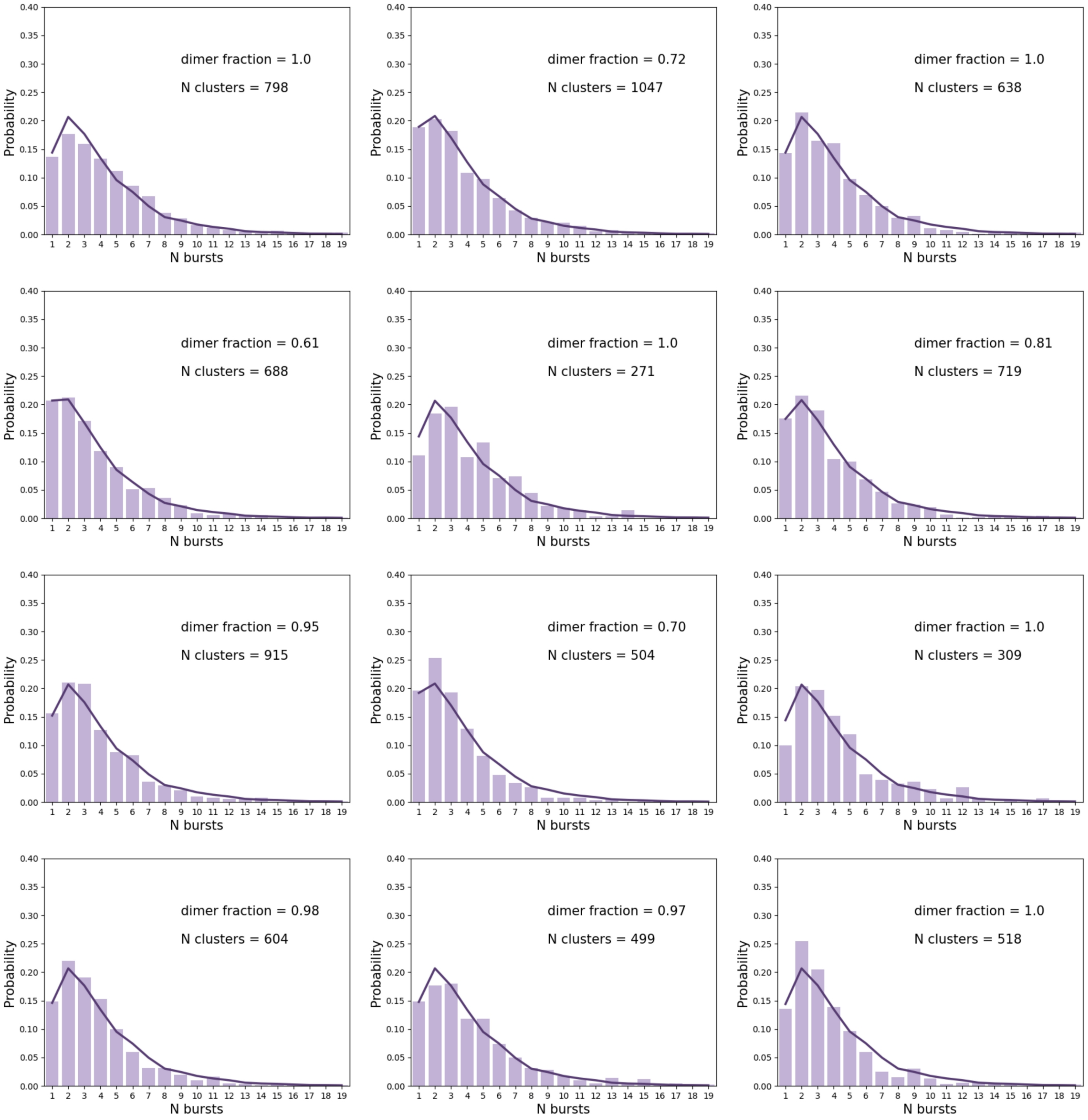

**Figure S4.** Burst distributions in clusters collected from individual cells expressing ChR2$_{C34A/C36A}$-mEos3.2. Each distribution was fitted with the weighted sum of reference monomeric and dimeric distributions, calculated dimer fraction is indicated.

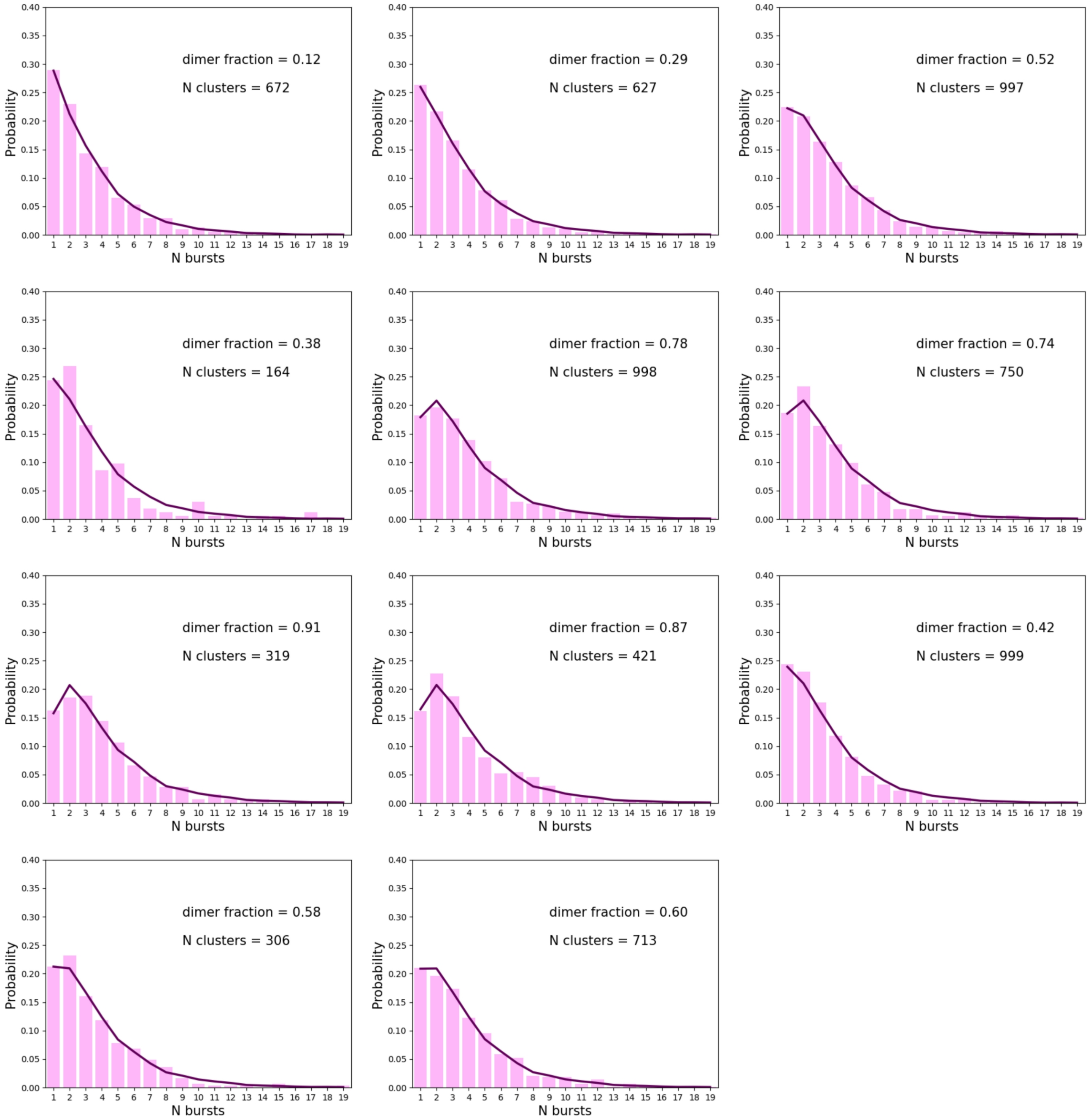

**Table S1.** DNA sequences encoding proteins used in this study (sequences of linkers between membrane protein and mEos3.2 genes are shown in lowercase)

| Name of the construct | DNA sequence |
|---|---|
| $\beta_1$AR-mEos3.2 | ATGGGCGCGGGGGTGCTCGTCCTGGGCGCCTCCGAGCCCGGT<br>AACCTGTCGTCGGCCGCACCGCTCCCCGACGGCGCGGCCACCG<br>CGGCGCGGCTGCTGGTGCCCGCGTCGCCGCCCGCCTCGTTGCT |

| | |
|---|---|
| | GCCTCCCGCCAGCGAAAGCCCCGAGCCGCTGTCTCAGCAGTGG<br>ACAGCGGGCATGGGTCTGCTGATGGCGCTCATCGTGCTGCTCAT<br>CGTGGCGGGCAATGTGCTGGTGATCGTGGCCATCGCCAAGACG<br>CCGCGGCTGCAGACGCTCACCAACCTCTTCATCATGTCCCTGGC<br>CAGCGCCGACCTGGTCATGGGGCTGCTGGTGGTGCCGTTCGGG<br>GCCACCATCGTGGTGTGGGGCCGCTGGGAGTACGGCTCCTTCT<br>TCTGCGAGCTGTGGACCTCAGTGGACGTGCTGTGCGTGACGGC<br>CAGCATCGAGACCCTGTGTGTCATTGCCCTGGACCGCTACCTCG<br>CCATCACCTCGCCCTTCCGCTACCAGAGCCTGCTGACGCGCGC<br>GCGGGCGCGGGGCCTCGTGTGCACCGTGTGGGCCATCTCGGC<br>CCTGGTGTCCTTCCTGCCCATCCTCATGCACTGGTGGCGGGCG<br>GAGAGCGACGAGGCGCGCCGCTGCTACAACGACCCCAAGTGCT<br>GCGACTTCGTCACCAACCGGGCCTACGCCATCGCCTCGTCCGTA<br>GTCTCCTTCTACGTGCCCCTGTGCATCATGGCCTTCGTGTACCT<br>GCGGGTGTTCCGCGAGGCCCAGAAGCAGGTGAAGAAGATCGAC<br>AGCTGCGAGCGCCGTTTCCTCGGCGGCCCAGCGCGGCCGCCCT<br>CGCCCTCGCCCTCGCCCGTCCCCGCGCCCGCGCCGCCGCCCG<br>GACCCCCGCGCCCCGCCGCCGCCGCCGCCACCGCCCCGCTGG<br>CCAACGGGCGTGCGGGTAAGCGGCGGCCCTCGCGCCTCGTGG<br>CCCTGCGCGAGCAGAAGGCGCTCAAGACGCTGGGCATCATCAT<br>GGGCGTCTTCACGCTCTGCTGGCTGCCCTTCTTCCTGGCCAACG<br>TGGTGAAGGCCTTCCACCGCGAGCTGGTGCCCGACCGCCTCTT<br>CGTCTTCTTCAACTGGCTGGGCTACGCCAACTCGGCCTTCAACC<br>CCATCATCTACTGCCGCAGCCCCGACTTCCGCAAGGCCTTCCAG<br>GGACTGCTCTGCTGCGCGCGCAGGGCTGCCCGCCGGCGCCAC<br>GCGACCCACGGAGACCGGCCGCGCGCCTCGGGCTGTCTGGCC<br>CGGCCCGGACCCCCGCCATCGCCCGGGGCCGCCTCGGACGAC<br>GACGACGACGATGTCGTCGGGGCCACGCCGCCCGCGCGCCTG<br>CTGGAGCCCTGGGCCGGCTGCAACGGCGGGGCGGCGGCGGAC<br>AGCGACTCGAGCCTGGACGAGCCGTGCCGCCCCGGCTTCGCCT<br>CGGAATCCAAGGTGtctagaATGAGTGCGATTAAGCCAGACATGAA<br>GATCAAACTCCGTATGGAAGGCAACGTAAACGGGCACCACTTTG<br>TGATCGACGGAGATGGTACAGGCAAGCCTTTTGAGGGAAAACAG<br>AGTATGGATCTTGAAGTCAAAGAGGGCGGACCTCTGCCTTTTGC<br>CTTTGATATCCTGACCACTGCATTCCATTACGGCAACAGGGTATT<br>CGCCAAATATCCAGACAACATACAAGACTATTTTAAGCAGTCGTT<br>TCCTAAGGGGTATTCGTGGGAACGAAGCTTGACTTTCGAAGACG<br>GGGGCATTTGCAACGCCAGAAACGACATAACAATGGAAGGGGA<br>CACTTTCTATAATAAAGTTCGATTTTATGGTACCAACTTTCCCGCC<br>AATGGTCCAGTTATGCAGAAGAAGACGCTGAAATGGGAGCCCTC<br>CACTGAGAAAATGTATGTGCGTGATGGAGTGCTGACGGGTGATA<br>TTGAGATGGCTTTGTTGCTTGAAGGAAATGCCCATTACCGATGTG<br>ACTTCAGAACTACTTACAAAGCTAAGGAGAAGGGTGTCAAGTTAC<br>CAGGCGCCCACTTTGTGGACCACTGCATTGAGATTTTAAGCCAT<br>GACAAAGATTACAACAAGGTTAAGCTGTATGAGCATGCTGTTGCT<br>CATTCTGGATTGCCTGACAATGCCAGACGAAGATAA |
| CD28-mEos3.2 | ATGCTCAGGCTGCTCTTGGCTCTCAACTTATTCCCTTCAATTCAA<br>GTAACAGGAAACAAGATTTTGGTGAAGCAGTCGCCCATGCTTGT<br>AGCGTACGACAATGCGGTCAACCTTAGCTGCAAGTATTCCTACA<br>ATCTCTTCTCAAGGGAGTTCCGGGCATCCCTTCACAAAGGACTG<br>GATAGTGCTGTGGAAGTCTGTGTTGTATATGGGAATTACTCCCAG<br>CAGCTTCAGGTTTACTCAAAAACGGGGTTCAACTGTGATGGGAA<br>ATTGGGCAATGAATCAGTGACATTCTACCTCCAGAATTTGTATGT<br>TAACCAAACAGATATTTACTTCTGCAAAATTGAAGTTATGTATCCT<br>CCTCCTTACCTAGACAATGAGAAGAGCAATGGAACCATTATCCAT<br>GTGAAAGGGAAACACCTTTGTCCAAGTCCCCTATTTCCCGGACC |

| | |
|---|---|
| | TTCTAAGCCCTTTTGGGTGCTGGTGGTGGTTGGTGGAGTCCTGG<br>CTTGCTATAGCTTGCTAGTAACAGTGGCCTTTATTATTTCTGGG<br>TGAGGAGTAAGAGGAGCAGGccaggaggagcggccgccATGAGTGCGA<br>TTAAGCCAGACATGAAGATCAAACTCCGTATGGAAGGCAACGTA<br>AACGGGCACCACTTTGTGATCGACGGAGATGGTACAGGCAAGC<br>CTTTTGAGGGAAAACAGAGTATGGATCTTGAAGTCAAAGAGGGC<br>GGACCTCTGCCTTTTGCCTTTGATATCCTGACCACTGCATTCCAT<br>TACGGCAACAGGGTATTCGCCAAATATCCAGACAACATACAAGA<br>CTATTTTAAGCAGTCGTTTCCTAAGGGGTATTCGTGGGAACGAAG<br>CTTGACTTTCGAAGACGGGGGCATTTGCAACGCCAGAAACGACA<br>TAACAATGGAAGGGGACACTTTCTATAATAAAGTTCGATTTTATG<br>GTACCAACTTTCCCGCCAATGGTCCAGTTATGCAGAAGAAGACG<br>CTGAAATGGGAGCCCTCCACTGAGAAAATGTATGTGCGTGATGG<br>AGTGCTGACGGGTGATATTGAGATGGCTTTGTTGCTTGAAGGAA<br>ATGCCCATTACCGATGTGACTTCAGAACTACTTACAAAGCTAAGG<br>AGAAGGGTGTCAAGTTACCAGGCGCCCACTTTGTGGACCACTGC<br>ATTGAGATTTTAAGCCATGACAAAGATTACAACAAGGTTAAGCTG<br>TATGAGCATGCTGTTGCTCATTCTGGATTGCCTGACAATGCCAGA<br>CGAAGATAA |
| ChR2$_{WT}$-mEos3.2 | ATGGACTATGGCGGCGCTTTGTCTGCCGTCGGACGCGAACTTTT<br>GTTCGTTACTAATCCTGTGGTGGTGAACGGGTCCGTCCTGGTCC<br>CTGAGGATCAATGTTACTGTGCCGGATGGATTGAATCTCGCGGC<br>ACGAACGGCGCTCAGACCGCGTCAAATGTCCTGCAGTGGCTTG<br>CAGCAGGATTCAGCATTTTGCTGCTGATGTTCTATGCCTACCAAA<br>CCTGGAAATCTACATGCGGCTGGGAGGAAATCTATGTGTGCGCC<br>ATTGAAATGGTTAAGGTGATTCTGGAGTTCTTTTTTGAGTTTAAGA<br>ATCCCTCTATGCTCTACCTTGCCACAGGACACCGGGTGCAGTGG<br>CTGCGCTATGCAACCTGGCTGCTCACTTGTCCTGTCATCCTTATC<br>CACCTGAGCAACCTCACCGGCCTGAGCAACGACTACAGCAGGA<br>GAACCATGGGACTCCTTGTCTCAGACATCGGGTGCATCGTGTGG<br>GGGGCTACCAGCGCCATGGCAACCGGCTATGTTAAAGTCATCTT<br>CTTTTGTCTTGGATTGTGCTATGGCGCGAACACATTTTTTCACGC<br>CGCCAAAGCATATATCGAGGGTTATCATACTGTGCCAAAGGGTC<br>GGTGCCGCCAGGTCGTGACCGGCATGGCATGGCTGTTTTTCGT<br>GAGCTGGGGTATGTTCCCAATTCTCTTCATTTTGGGGCCCGAAG<br>GTTTTGGCGTCCTGAGCGTCTATGGCTCCACCGTAGGTCACACG<br>ATTATTGATCTGATGAGTAAAAATTGTTGGGGGTTGTTGGGACAC<br>TACCTGCGCGTCCTGATCCACGAGCACATATTGATTCACGGAGA<br>TATCCGCAAAACCACCAAACTGAACATCGGCGGAACGGAGATCG<br>AGGTCGAGACTCTCGTCGAAGACGAAGCCGAGGCCGGAGCCGT<br>GccagcggccgccATGAGTGCGATTAAGCCAGACATGAAGATCAAAC<br>TCCGTATGGAAGGCAACGTAAACGGGCACCACTTTGTGATCGAC<br>GGAGATGGTACAGGCAAGCCTTTTGAGGGAAAACAGAGTATGGA<br>TCTTGAAGTCAAAGAGGGCGGACCTCTGCCTTTTGCCTTTGATAT<br>CCTGACCACTGCATTCCATTACGGCAACAGGGTATTCGCCAAAT<br>ATCCAGACAACATACAAGACTATTTTAAGCAGTCGTTTCCTAAGG<br>GGTATTCGTGGGAACGAAGCTTGACTTTCGAAGACGGGGGCATT<br>TGCAACGCCAGAAACGACATAACAATGGAAGGGGACACTTTCTA<br>TAATAAAGTTCGATTTTATGGTACCAACTTTCCCGCCAATGGTCC<br>AGTTATGCAGAAGAAGACGCTGAAATGGGAGCCCTCCACTGAGA<br>AAATGTATGTGCGTGATGGAGTGCTGACGGGTGATATTGAGATG<br>GCTTTGTTGCTTGAAGGAAATGCCCATTACCGATGTGACTTCAGA<br>ACTACTTACAAAGCTAAGGAGAAGGGTGTCAAGTTACCAGGCGC<br>CCACTTTGTGGACCACTGCATTGAGATTTTAAGCCATGACAAAGA<br>TTACAACAAGGTTAAGCTGTATGAGCATGCTGTTGCTCATTCTGG<br>ATTGCCTGACAATGCCAGACGAAGATAA |

| | |
|---|---|
| ChR2$_{C34A/C36A}$-mEos3.2 | ATGGACTATGGCGGCGCTTTGTCTGCCGTCGGACGCGAACTTTT<br>GTTCGTTACTAATCCTGTGGTGGTGAACGGGTCCGTCCTGGTCC<br>CTGAGGATCAAGCTTACGCTGCCGGATGGATTGAATCTCGCGGC<br>ACGAACGGCGCTCAGACCGCGTCAAATGTCCTGCAGTGGCTTG<br>CAGCAGGATTCAGCATTTTGCTGCTGATGTTCTATGCCTACCAAA<br>CCTGGAAATCTACATGCGGCTGGGAGGAAATCTATGTGTGCGCC<br>ATTGAAATGGTTAAGGTGATTCTGGAGTTCTTTTTTGAGTTTAAGA<br>ATCCCTCTATGCTCTACCTTGCCACAGGACACCGGGTGCAGTGG<br>CTGCGCTATGCAACCTGGCTGCTCACTTGTCCTGTCATCCTTATC<br>CACCTGAGCAACCTCACCGGCCTGAGCAACGACTACAGCAGGA<br>GAACCATGGGACTCCTTGTCTCAGACATCGGGTGCATCGTGTGG<br>GGGGCTACCAGCGCCATGGCAACCGGCTATGTTAAAGTCATCTT<br>CTTTTGTCTTGGATTGTGCTATGGCGCGAACACATTTTTTCACGC<br>CGCCAAAGCATATATCGAGGGTTATCATACTGTGCCAAAGGGTC<br>GGTGCCGCCAGGTCGTGACCGGCATGGCATGGCTGTTTTTCGT<br>GAGCTGGGGTATGTTCCCAATTCTCTTCATTTTGGGGCCCGAAG<br>GTTTTGGCGTCCTGAGCGTCTATGGCTCCACCGTAGGTCACACG<br>ATTATTGATCTGATGAGTAAAAATTGTTGGGGGTTGTTGGGACAC<br>TACCTGCGCGTCCTGATCCACGAGCACATATTGATTCACGGAGA<br>TATCCGCAAAACCACCAAACTGAACATCGGCGGAACGGAGATCG<br>AGGTCGAGACTCTCGTCGAAGACGAAGCCGAGGCCGGAGCCGT<br>GccagcggccgccATGAGTGCGATTAAGCCAGACATGAAGATCAAAC<br>TCCGTATGGAAGGCAACGTAAACGGGCACCACTTTGTGATCGAC<br>GGAGATGGTACAGGCAAGCCTTTTGAGGGAAAACAGAGTATGGA<br>TCTTGAAGTCAAAGAGGGCGGACCTCTGCCTTTTGCCTTTGATAT<br>CCTGACCACTGCATTCCATTACGGCAACAGGGTATTCGCCAAAT<br>ATCCAGACAACATACAAGACTATTTTAAGCAGTCGTTTCCTAAGG<br>GGTATTCGTGGGAACGAAGCTTGACTTTCGAAGACGGGGGCATT<br>TGCAACGCCAGAAACGACATAACAATGGAAGGGGACACTTTCTA<br>TAATAAAGTTCGATTTTATGGTACCAACTTTCCCGCCAATGGTCC<br>AGTTATGCAGAAGAAGACGCTGAAATGGGAGCCCTCCACTGAGA<br>AAATGTATGTGCGTGATGGAGTGCTGACGGGTGATATTGAGATG<br>GCTTTGTTGCTTGAAGGAAATGCCCATTACCGATGTGACTTCAGA<br>ACTACTTACAAAGCTAAGGAGAAGGGTGTCAAGTTACCAGGCGC<br>CCACTTTGTGGACCACTGCATTGAGATTTTAAGCCATGACAAAGA<br>TTACAACAAGGTTAAGCTGTATGAGCATGCTGTTGCTCATTCTGG<br>ATTGCCTGACAATGCCAGACGAAGATAA |
